# Gene deletion as a possible strategy adopted by New World *Leishmania infantum* to maximize geographic dispersion

**DOI:** 10.1101/2024.05.22.595165

**Authors:** Monique Florêncio, Marne Coimbra Chagas, Anderson Guimarães-Costa, Jullyanna Oliveira, Ingrid Waclawiak, Thamara K. F. Oliveira, Elvira Maria Saraiva, Anita Leocadio Freitas-Mesquita, José Roberto Meyer-Fernandes, Laura Aragão-Farias, Camilly Enes Trindade, Patricia Cuervo Escobar, Renata Azevedo do Nascimento, Otacilio C. Moreira, Flávia Lima Ribeiro-Gomes, Yara M. Traub-Csekö, Erich Loza Telleria, Slavica Vaselek, Tereza Leštinová, Petr Volf, Gerald F. Späth, Elisa Cupolillo, Mariana C. Boité

**Affiliations:** Laboratório de Pesquisa em Leishmaniose, Instituto Oswaldo Cruz, FIOCRUZ, 21040-360 Rio de Janeiro, Brazil; Laboratório de Imunologia das Leishmanioses, Instituto de Microbiologia Paulo de Góes, Centro de Ciências da Saúde, Universidade Federal do Rio de Janeiro - UFRJ, Rio de Janeiro 21941-902, Brazil.; Instituto de Bioquímica Médica Leopoldo de Meis - IBqM, Universidade Federal do Rio de Janeiro - UFRJ, 21941-590. Rio de Janeiro, RJ, Brazil; Laboratory of Molecular Virology and Parasitology, Instituto Oswaldo Cruz, FIOCRUZ, Rio de Janeiro, RJ, Brazil. 21040-360, Brazil; Laboratório de Pesquisa em Malaria, Instituto Oswaldo Cruz, FIOCRUZ, Rio de Janeiro, RJ, Brazil; Laboratório de Biologia Molecular de Parasitas e Vetores, Instituto Oswaldo Cruz, 21040-365 Rio de Janeiro, Brazil; Department of Parasitology, Faculty of Science, Charles University, 128 44 Prague, Czech Republic; Institut Pasteur, Univerisité Paris Cité, INSERM U1201, Unité de Parasitologie moléculaire et Signalisation, 75015 Paris, France.; Instituto Nacional de Ciência e Tecnologia EpiAMO (INCT)

**Author notes:** **Author’s contribution: Conceptualization** GS, EC, MCB. **Methodology**, MF, MCC, AG-C, JO, IW, TO, ALF-M, LA, CE, RAN, FLR-G, ELT, SV, TL, MCB. **Formal analysis** MF, AGC, OCM, ELT, MCB. **Data curation** MF, MCC, AGC, ALF-M, OCM, ELT, MCB. **Writing—original draft preparation** MF, MCB. **Writing—review and editing** AG-C, EMS, ALF-M, JRM-F, PC, OCM, FLR- G, YMT-C, ELT, PV, GS, EC. **Visualization** MF, AGC, ELT, MCB. **Supervision** EMS, JRM-F, PC, YMT-C, PV, MCB. **Project administration** MCB. **Funding acquisition** EMS, AGC, JRM-F, YMT-C, PV, GS, EC, MCB.

**Keywords:** *Leishmania infantum*, visceral leishmaniasis, Brazil, ectonucleotidase, parasite fitness, host-parasite, *Leishmania* transmission

## Abstract

**Background:** The present study investigates implications of a sub-chromosomal deletion in *Leishmania infantum* strains, the causative agent of American Visceral Leishmaniasis (AVL). Primarily found in New World strains, the deletion leads to the absence of the ecto-3’-nucleotidase/nuclease enzyme (3’NU/NT), impacting parasite virulence, pathogenicity, and drug susceptibility. The potential factors favoring prevalence and the widespread geographic distribution of these deleted mutant parasites (DEL) in the New World (NW) are discussed under the generated data.

**Methods:** We conducted phenotypic analyses of the parasites showing the sub-chromosomal deletion by applying *in vitro* assays of 3’NU/NT activity, metacyclic enrichment, and relative quantitation of transcripts abundance on axenic parasites. We further performed experimental infections in both *in vitro* and *in vivo* models of vertebrate and invertebrate hosts using geographically diverse mutant field isolates.

**Results:** Virulence assays, poorer ability to survive neutrophil traps (NETs) and murine model infection revealed reduced pathogenicity in vertebrate hosts by the DEL strains. Conversely, these parasites exhibit enhanced metacyclogenesis and colonization rates in sand flies, potentially facilitating transmission. This combination may represent a more efficient way to maintain and disperse the transmission cycle of DEL strains.

**Conclusions:** Phenotypic assessments reveal altered parasite fitness, with enhanced transmissibility at the population level. Reduced susceptibility of DEL strains to miltefosine, a key drug in VL treatment, further complicates control efforts. Our study underscores the importance of typing parasite genomes for surveillance and control and proposes the sub-chromosomal deletion as a molecular marker in AVL management.

## Introduction

*L. infantum* is one of the causative agents of Visceral Leishmaniasis (VL) across Asia, Africa, and Europe, and is exclusively responsible for American Visceral Leishmaniasis (AVL), affecting both humans and dogs, the primary urban reservoir. The course of this parasite’s evolution, adaptation and geographic dispersion is constantly under the effect of multivariable interactions in the transmission cycle, which include vertebrate hosts and various sand fly species as vectors. Within this process, the parasite’s intrinsic genome instability has been predicted as a driver in *Leishmania* fitness gain in response to environmental changes^1^ or drug pressure^2^. The loss of genes, for instance, is a molecular strategy eventually compensated by adjustments at the genomic, transcriptomic, and post-transcriptomic levels, leading to phenotypic variances^3^ that may favor parasite survival and transmission. In this context, our previous study has documented the prevalent distribution of New World (NW) *L. infantum* strains harboring a 12 Kb sub-chromosomal deletion (DEL mutant strains)^4^.

It is noteworthy that such deletions are frequently observed among analyzed genomes from Brazilian strains, primarily confined to parasite populations in the Americas^4^. This genomic trait promotes an important phenotypic change for the parasite, notably the absence of the ecto-3’-nucleotidase/nuclease (3’NU/NT) activity^4^, encoded by two of the four genes removed by the deletion. The 3’NU/NT is unique to trypanosomatids^5^, and, as an ecto- enzyme, acts upon substrates present extracellularly. Its role encompasses the hydrolysis of extracellular 3ʹ nucleotides and nucleic acids, generating nucleosides that can be translocated across plasma membranes via nucleoside transporters (NT)^5^. The 3’NU/NT enzyme is involved in the purine salvage pathway of *Leishmania*^6,7^ (trypanosomatids are unable to synthetize purine de novo^6^), thereby influencing nutrition^8^and parasite replication. Moreover, it is implicated as a virulence factor affecting the parasite’s ability to infect macrophages^9–11^ and survive neutrophil extracellular traps (NETs)^12,13^. Recent evidence has linked 3’NU/NT to susceptibility to miltefosine (MIL) ^14^, a critical drug in leishmaniasis treatment, and clinical trials have shown a correlation between reduced MIL efficacy in AVL treatment and infection by DEL strains^15^. Despite this, MIL is approved to be utilized in the treatment of canine visceral leishmaniasis (CVL), prompting concerns regarding surveillance and control^16^.

The prevalence and geographical spread of DEL strains in Brazil underscore a paradoxical hypothesis suggesting that the loss of two 3’NU/NT genes confers fitness advantages, facilitating the spread of the mutant population. This puzzling association raises fundamental questions regarding how the loss of an important virulence and biological function contributes to the high prevalence of the mutant DEL strains. We postulate that the genetic and biological differences revealed so far between DEL and non-deleted strains (NonDEL) alter transmissibility of these parasite genotypes by affecting colonization parameters in the vector, for instance, and/or resulting in variable infection outcomes in vertebrate hosts. To investigate this hypothesis, we performed phenotypic evaluations of parasites with the sub-chromosomal deletion. This involved conducting in vitro assays on axenic parasites and carrying out experimental infections in both in vitro and in vivo models, encompassing vertebrate and invertebrate hosts, using geographically diverse NonDEL and DEL field isolates.

Our findings reveal that DEL mutant parasites exhibit enhanced metacyclogenesis *in vitro* but diminished capacity to survive neutrophil NETs and infect macrophages. Although the uptake by macrophages of DEL parasites is reduced, the intracellular development and proliferation was not affected. *In vivo* analyses of selected DEL strains demonstrate often reduced recruitment of monocytes/neutrophils in a murine ear infection model, irrespective of strain origin. Moreover, colonization parameters in different vector species exhibit notable variances among strains from Central-West Brazil, underscoring the significant influence of sand flies on parasite distribution and indicating a fitness advantage for the intravectorial life stage of DEL strains.

Collectively, these data present a spectrum of phenotypes that, while individually possibly diminishing DEL parasite fitness, collectively contribute to enhanced transmissibility, as metacyclic forms are the infective stage transmitted by sand flies, and reduced pathogenicity in the vertebrate host may facilitate transmission for some parasites^17^. Additionally, the elevated colonization rate observed for DEL strains on key vector species enhances the likelihood of prevalence among these mutants in the Americas.

Further epidemiological implications related to the co-circulation of these *L. infantum* populations for VL treatment and control is the reduced susceptibility to MIL observed for DEL parasites^15,18^. Combined to the altered pathogenicity and infectivity described herein, the data emphasize the importance of detecting asymptomatic individuals and typing the infecting parasite’s genome as DEL or NonDEL, particularly for canine leishmaniasis. Consequently, we advocate for the sub-chromosomal deletion as a key molecular marker for VL surveillance in the American continent.

## Results

### Deletion-carrying strains present relevant biological differences

Initially, a set of 23 culture-adapted *L. infantum* strains, isolated from humans and dogs, comprising nine deletion-carrying (DEL), three heterozygous (HTZ), and eleven non-deleted (NonDEL) strains, were evaluated. Subsequently, subsets of strains were chosen for different assays, as outlined in the Methods section and Supp. material. Genotypes were identified via whole genome sequencing (WGS)^4^ and further confirmed by PCR.

The diminished or absent ecto-3’-nucleotidase (3’NT) activity in deletion-carrying strains (DEL and HTZ) was validated across the entire sample panel. Variable 3’NT activity was observed among NonDEL samples, with strains exhibiting either higher, lower, or equivalent activity compared to a *L. amazonensis* reference strain (Figure 1A). As anticipated, 3’NT activity among heterozygous hybrids (HTZ) was below the average observed in NonDEL samples but higher than in the DEL group (Figure 1B). The intermediate 3’NT activity for HTZ reflects the impact of the intermediate copy number of the 3’NU/NT, as previously confirmed by qPCR^4^, and an intermediate read depth within the deleted site, as confirmed by WGS.

**Figure 1.**
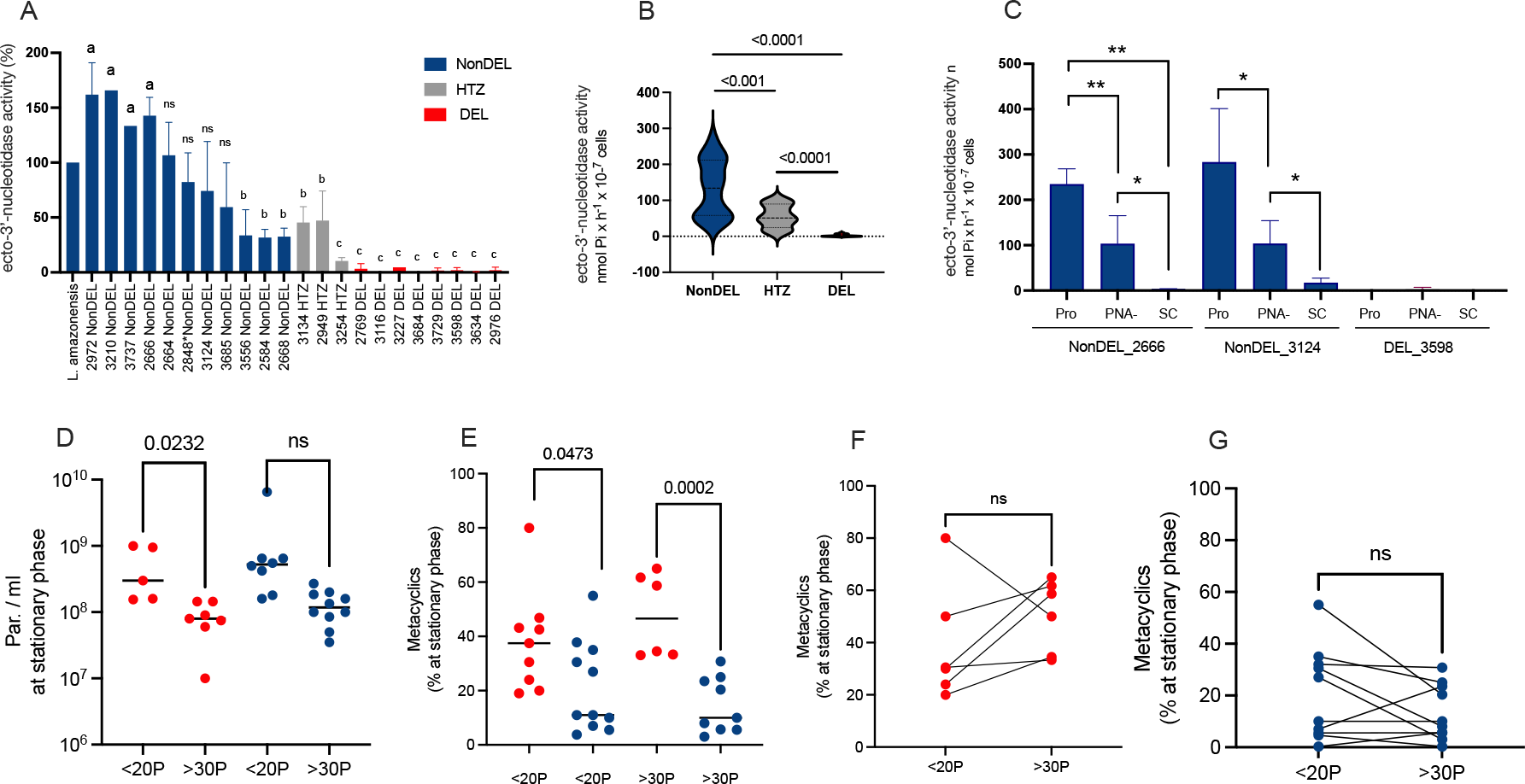
In vitro phenotypic characterization of DEL, NonDEL and HTZ strains. A) Normalized 3’NT activity for NonDEL, HTZ and DEL strains. The 3’NT activity was measured for DEL (n=08), HTZ (n=03) and NonDEL (n=11) parasites harvested at stationary phase during at least three independent experimental replicates. The original values, expressed as nmol Pi x h^-1^ x 10 ^-7^ cells, were normalized by the values obtained with the L. amazonensis assays. Compared to the L. amazonensis control, four NonDEL samples presented higher activity (a); three NonDEL and two HTZ strains presented lower activity (b); all DEL and one HTZ samples presented the lowest enzyme activity (c). Unpaired t-test *p<0.05; **p<0.01. B) 3’NT activity average expressed as nmol Pi x h^-1^ x 10 ^-7^ for each genotype group composed by the strains individually presented in graph A. ANOVA. 3’NT activity expressed as nmol Pi x h^-1^ x 10 ^-7^ cells in cultures from: stationary phase promastigotes = Pro; metacyclic enriched fraction of stationary phase culture = PNA -; SC = stressed culture parasites “axenic amastigotes”. D) Parasite density and E) percentage of metacyclic at stationary phase of culture with less than 20 passages (<20P) and adapted parasites - more than 30 passages (>30P). Percentage of metacyclic was determined 72 hours (late stationary phase) after an initial inoculum of 10^6^ parasites/ml. Metacyclic enrichment was obtained by Peanut agglutinin (PNA); percentage of cells from the PNA- fraction was determined in relation to the total cell count. Red = DEL; Blue=NonDEL; Grey=HTZ. Unpaired t test. F-G) Paired t-test for DEL and NonDEL groups of parasites in <20P and >30P conditions. ns=non-significant

A subset of three strains was selected for assays using promastigotes (Pro), PNA- enriched metacyclic, and parasites exposed to stress conditions (SC) encountered during amastigote differentiation (i.e. low pH and 37°C) to compare enzyme activity among these conditions. The highest 3’NT activity was observed in stationary phase NonDEL promastigote parasites (Pro; 237.7 and 283.1 nmol Pi x h-1 x 10^-7^ cells), followed by the metacyclic fraction (103.4 and 103.8 nmol Pi x h-1 x 10^-7^cells). These findings support previous reports indicating significant 3’NU/NT activity among metacyclic forms, suggesting the enzyme’s potential relevance during the intravectorial phase or early infection stages in the vertebrate host.

However, enzyme activity in tissue-derived amastigotes has not been measured to date. Notably, cultures exposed to pH/temperature stress (SC) displayed reduced or absent activity (3.3 and 17.4 nmol Pi x h-1 x 10^-7^ cells; **Figure 1**C). As expected, the DEL sample exhibited no activity across any lifecycle stage (Pro=0 and PNA- = 3.3 nmol Pi x h-1 x 10^-7^ cells) or in pH/temperature stress exposed cultures.

Low pH and nutritional stress, including the absence of purines^21,19^, trigger metacyclogenesis in trypanosomatides^19,20^. Trypanosomes and *Leishmania* lack a de novo purine biosynthetic pathway^6^, therefore, these parasites rely on alternative purine salvage routes involving enzymes such as the 3NU/NT. Reports^5,22^ suggest that the lack of 3’NT activity affects the parasite nutritional state and replication, thereby triggering metacyclogenesis. That information led us to test whether the parasites with distinct 3’NT activity levels differed in metacyclogenesis and cell density in vitro. Inoculum-controlled cultures (N=17 strains) with fewer than 20 passages (<20P) were submitted to metacyclic enrichment at the late stationary phase. Results revealed similar cell density, but higher percentage of metacyclic for DEL strains (median=58.6%) compared to NonDEL/HTZ (median=9%) strains (**Figure 1**D-E). To evaluate how consistent the difference in metacyclogenesis between groups was, the assay was repeated after 30 passages in culture (>30P) (**Figure 1**D-E). The difference was verified with statistical significance (p=0.002). Further paired analysis showed no notable variation in metacyclogenesis between cultures maintained for <20P and >30P (**Figure 1**F-G). Thus, despite the acknowledged impact of culture maintenance on *Leishmania* biology, the contrast in metacyclogenesis between DEL and NonDEL parasites remains consistent and robust.

### Higher abundance transcripts of Nucleoside Transporter 1 and META2 was detected among DEL strains

After searching for differential genomic signals between genotypes - besides the sub-chromosomal deletion itself, Schwabl et al.^4^ detected among DEL strains higher gene copy number of the NT1 adenosine/pyrimidine nucleoside transporter gene (NT1), involved in the salvage route for purines by promoting the uptake of extracellular adenosine by the parasite^23^. Moreover, starvation of *Leishmania donovani* parasites for purines leads to a rapid amplification of the abundance of transcripts for NT1^8^ gene. Based on these statements, the higher expression of this membrane transporter could represent a compensatory effect for the 3’NU/NT deficient parasites. In the same analysis^4^, the mutant strains showed reduced gene copy number for Paraflagellar and Amastin genes, which are associated with the parasite’s ability to infect mononuclear cells^24^. To assess whether these copy number variations (CNVs) are mirrored on transcript levels, we conducted relative quantitation of gene copy number using Real-Time PCR under two conditions: i) total RNA isolated from culture-adapted promastigotes harvested at the late stationary phase (**Figure 2**), and ii) total RNA isolated from metacyclic parasites enriched from late stationary phase culture (PNA-; **Figure 3**). The results confirmed the expected decrease in transcripts for 3’NU/NT among DEL samples in stationary-phase promastigotes (**Figure 2**A), but not in PNA- metacyclic parasites (**Figure 3A**). Additionally, the number of transcripts corroborated the CNV data for NT1^4^, with higher fold change in transcript abundance observed in the DEL group (2.0 vs 0.8 fold change median; **Figure 2**B). This suggests that alternative molecular pathways may emerge in mutant parasites. The results for Paraflagellar or Amastin transcripts (**Figure 2**C-D) did not align with previous CNV data, suggesting that a potential association between lower Paraflagellar and Amastin expression and reduced infectiousness in DEL samples is not supported.

**Figure 2.**
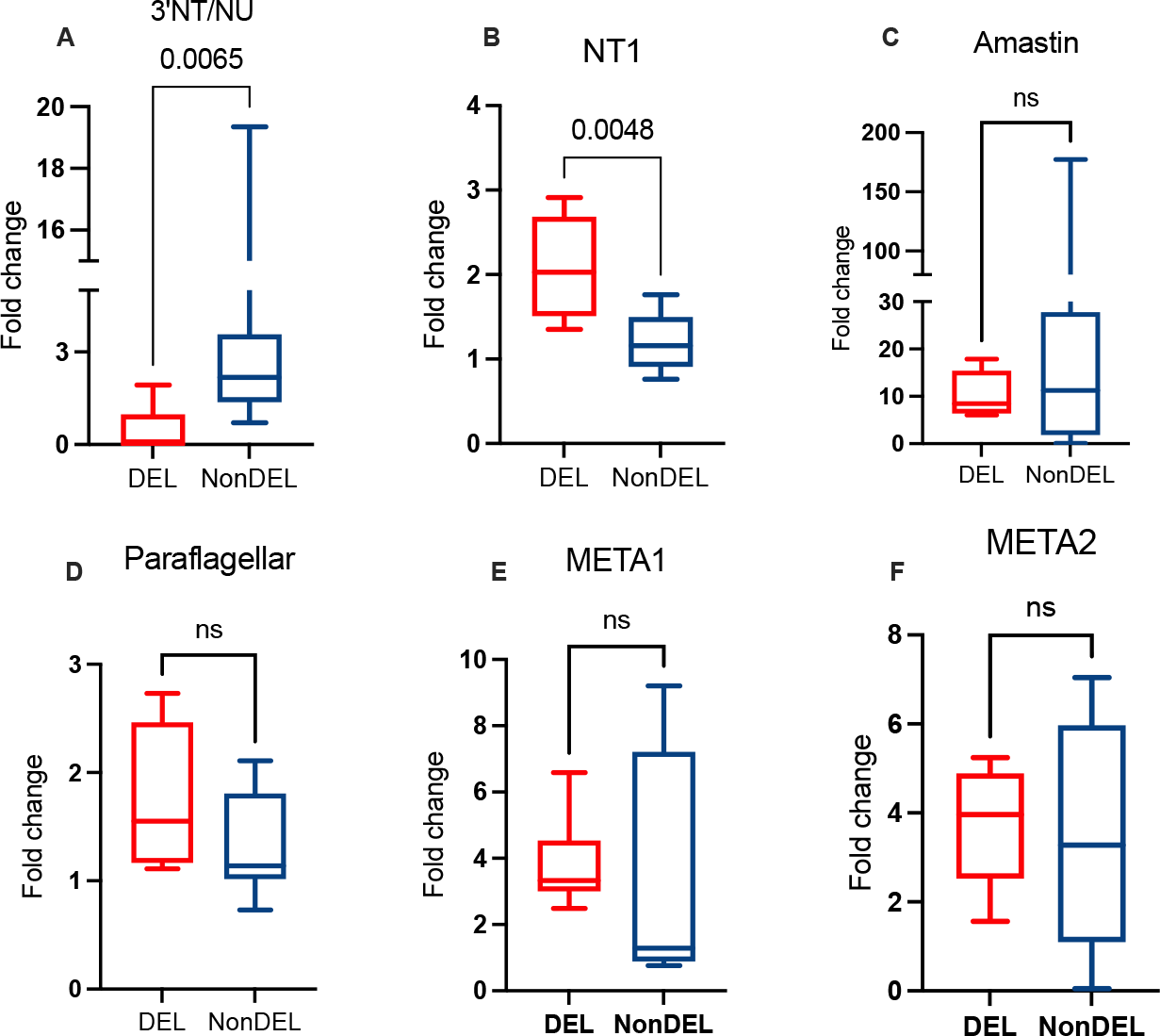
Fold change of transcripts from culture adapted promastigotes harvest from the late stationary phase (72 hours). The targets 3’NU/NT = 3’ecto-nucleotidase; NT1 = nucleoside transporter 1, Amastin and Paraflagellar were chosen based on previous data pointing to differences in CNV between DEL and NonDEL^4^. META1 and META2 were included as potential markers for metacyclic^25^ and for resistance to oxidative stress^26^. Total RNA was reversed transcribed in cDNA and targets quantified by Real Time qPCR. Delta-Delta Ct method was applied using alfa-tubulin as endogenous control and the OW sample NonDEL 3124 as calibrator. Mann Whitney t-test (unpaired) for DEL (n= 5) and NonDEL (n=9) samples. ns = no significant.

**Figure 3.**
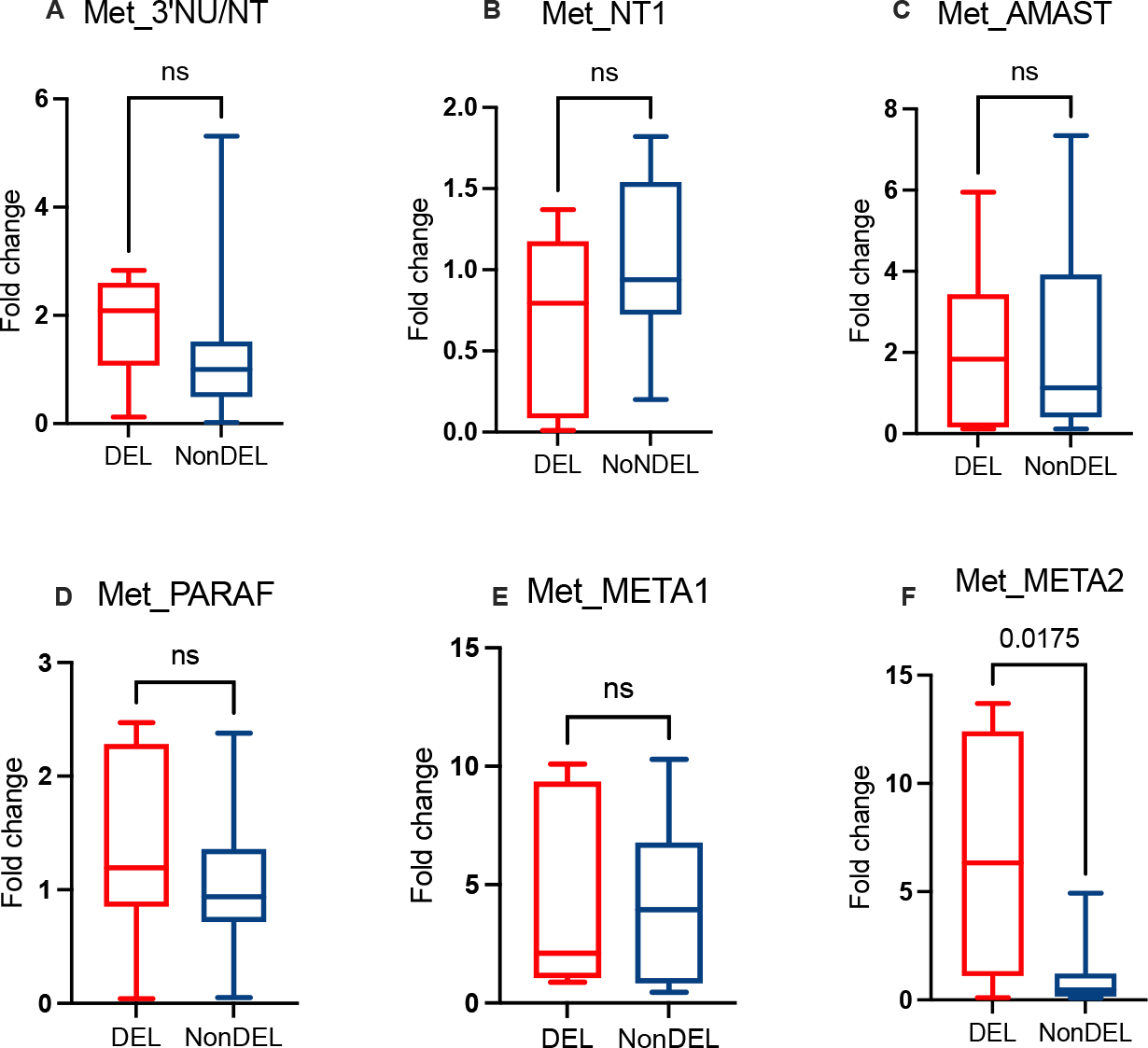
Fold change of transcripts from metacyclic-enriched cultures obtained by PNA from the late stationary phase (PNA-). The targets 3’NU/NT = 3’ecto-nucleotidase; NT1 = nucleoside transporter 1, Amastin and Paraflagellar were chosen based on previous data pointing to differences in CNV between DEL and NonDEL^4^. META1 and META2 were included as potential markers for metacyclic^25^ and for resistance to oxidative stress^26^. Total RNA from both PNA+ and PNA- cultures were reversed transcribed in cDNA and targets quantified by Real Time qPCR. Delta-Delta Ct method was applied using alfa- tubulin as reference gene and the sample in PNA+ used as a calibrator for the correspondent sample in PNA- culture. Mann Whitney Rank Sum Test or t-test (unpaired) for DEL (n= 5) and NonDEL (n=9) samples. ns = no significant.

META1 and META2 expression levels were assayed as potential markers for metacyclogenesis^25^ and for resistance to oxidative stress, as reported for *L. amazonensis*^26^. Both META1 and META2 presented similar transcript levels between DEL and NonDEL groups of strains. In PNA- samples, however (Figure 3), the DEL group exhibits higher fold change (6.6 vs 1.0; Figure 3F) for META2, suggesting these metacyclics may be more resistant to oxidative stress and heat shock^26^. No significative differences were detected for the other targets in PNA- cultures.

### Sand flies are differentially colonized by DEL and NonDEL strains from specific geographic areas

We assessed the impact of the main vector for visceral leishmaniasis in Brazil, *L. longipalpis*^27^, on the spread of DEL strains. Sand flies were infected with DEL from Rio de Janeiro (RJ) (N=1), NonDEL from Mato Grosso do Sul (MS) (N=1), or HTZ strains (N=1). After 192 hours, infection rates were similar, (Figure 4A) but DEL and HTZ strains showed higher parasite loads per midgut compared to NonDEL, indicating that the deletion potentially increases fitness during the vector life cycle stage.

**Figure 4.**
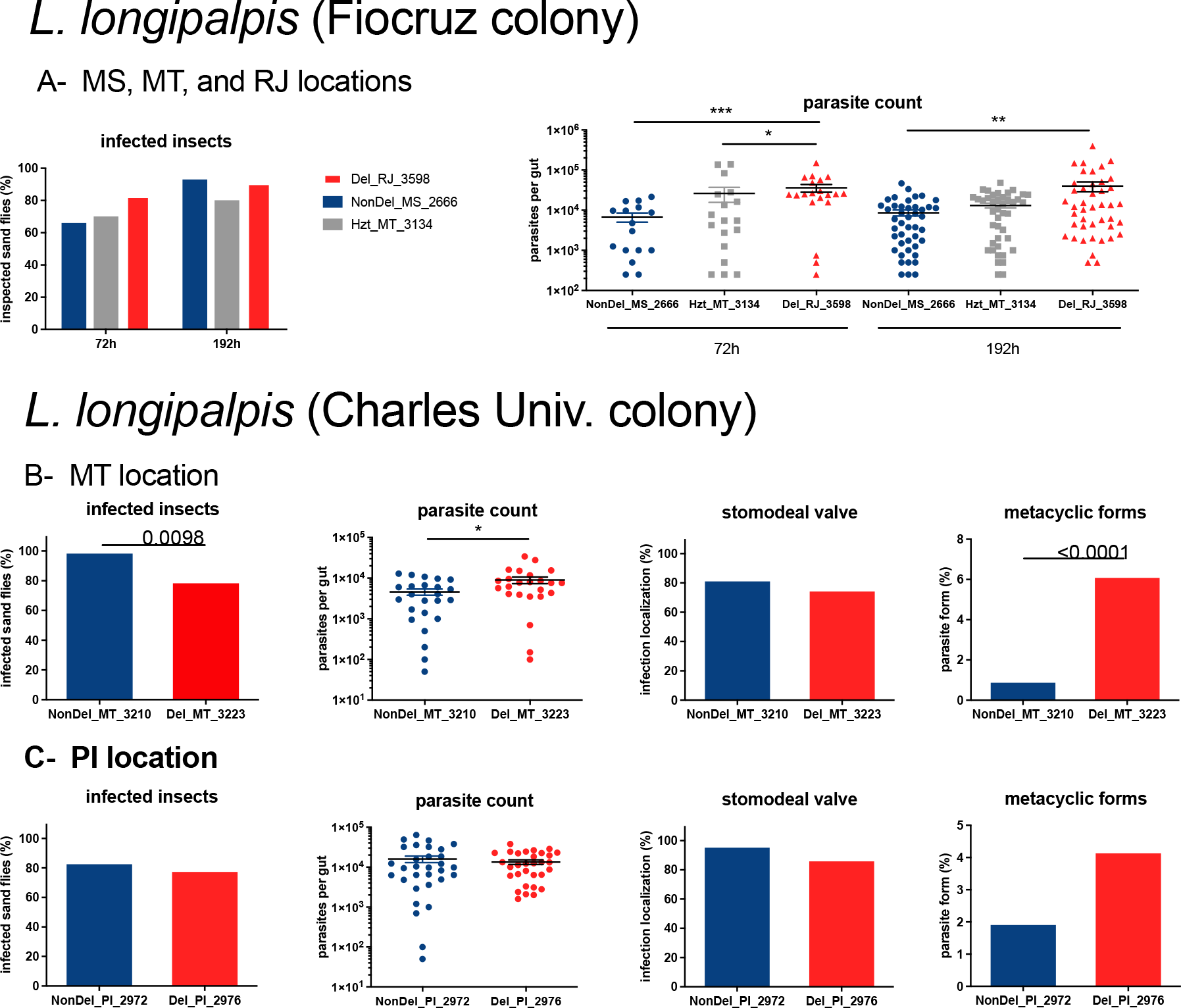
Infections with DEL and Non-DEL strains from specific geographic areas in L. longipalpis. – (**A**) Percentage of infected L. longipalpis and parasite number of NonDEL_MS_2666 (blue), HTZ_MT_3134 (green), and DEL_RJ_3598 (red) strains. **B** and **C**. Percentage of infected L. longipalpis, parasite number, percentage of insects that developed infection to the stomodeal valve, and percentage of metacyclic parasite forms of (**B**) NonDEL_MT_3210 (blue) and DEL_MT_3223 (red) strains from MT location, and (**C**) NonDEL_PI_2972 (blue) and DEL_PI_2976 (red) strains from PI location. Statistically significant differences: *p < 0.05, ** p < 0.01, ***p<0.001 and ****p < 0.0001.

To explore the influence of strain geographic origin on *L. longipalpis* and *L. migonei* infections additional DEL (n=2) and NonDEL (n=2) samples from Mato Grosso (MT), and Piauí (PI) were included. In *L. longipalpis*, the DEL strain from MT exhibited similar behavior to the MS/RJ DEL, with higher parasite counts after 192 hours despite a lower number of infected insects. Moreover, percentages of metacyclic forms were increased for DEL strains (Figure 4B). In contrast, no significant differences were observed for the second pair of DEL and NonDEL *Leishmania* from PI (Figure 4C), but, again, an increased percentage of metacyclic forms was observed in DEL strain. These findings suggest a potential link between vector species and the spread of specific parasite strains in Brazilian conditions.

To assess the adaptation of parasites to different sand fly species, we examined the impact of the sub-chromosomal deletion during infections in *L. migonei,* a sand fly species involved in parasite dispersion in some locations in Brazil, and *P. perniciosus* an Old World sand fly vector of *L. infantum*. In *L. migonei*, no significant differences were found in infection parameters, though a slight increase in stomodeal valve colonization and metacyclic form numbers was noted with the DEL strain from the MT location (**Figure 5**A). A similar trend was observed in *P. perniciosus*, where significant differences were observed (**Figure 5**C-D).

**Figure 5.**
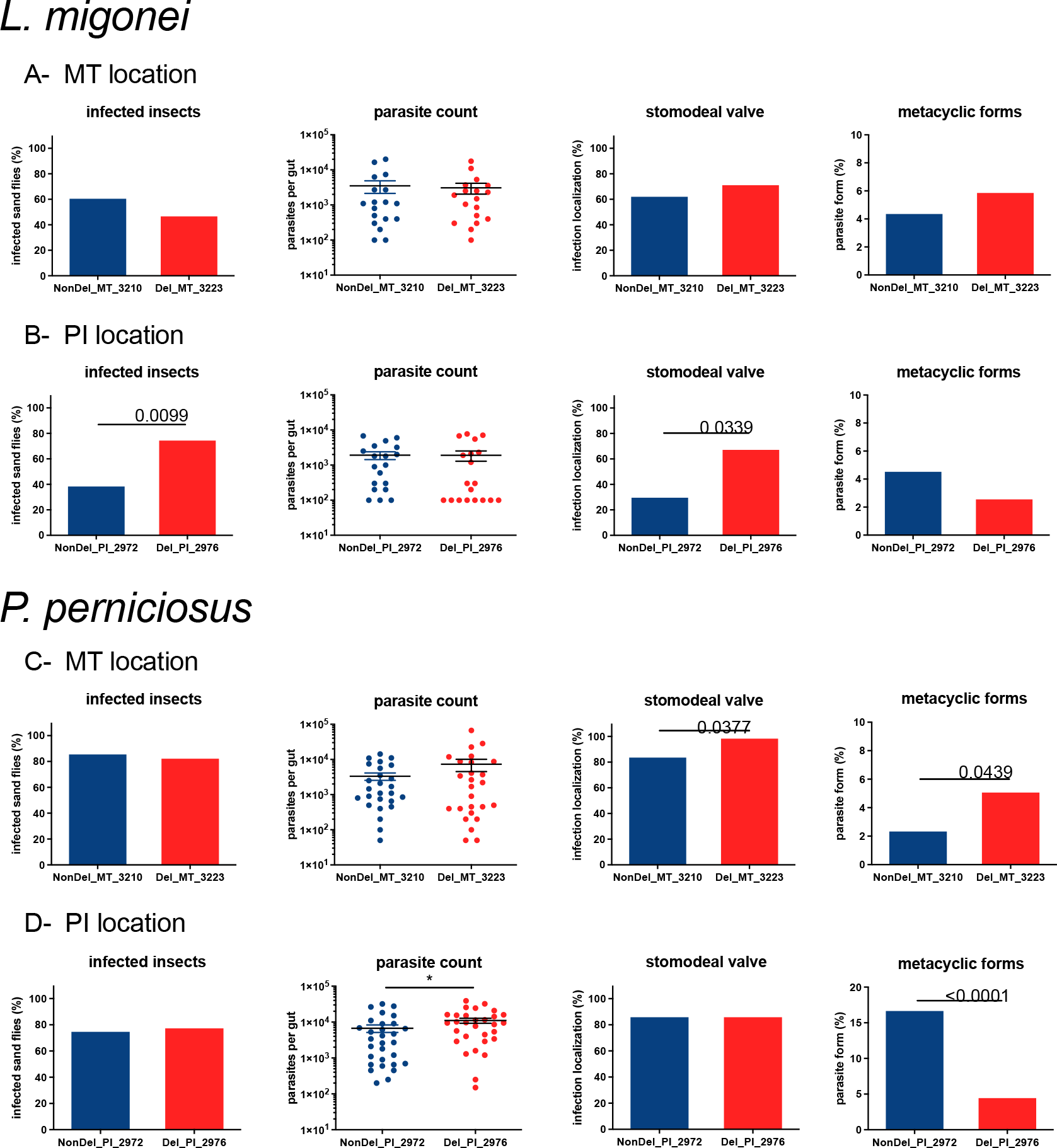
Infections with DEL and Non-DEL strains from specific geographic areas in L. migonei and P. perniciosus. – Percentage of infected (**A** and **B**) L. migonei and (**C** and **D**) P. perniciosus, parasite number, percentage of insects that developed infection to the stomodeal valve, and percentage of metacyclic parasite forms of (**A** and **B**) NonDEL_MT_3210 (blue) and DEL_MT_3223 (red) strains from MT location, and (**C** and **D**) NonDEL_PI_2972 (blue) and DEL_PI_2976 (red) strains from PI location. .

For strains from PI, the percentage of *L. migonei* sand flies infected and colonized stomodeal valve was increased in DEL strains fed group (76% vs 40% in NonDEL; 68.7% vs 31.25 in NonDEL, respectively), but with slightly fewer metacyclic parasites (2.7% vs 4.6% in NonDEL) (**Figure 5**B). In *P. perniciosus*, the same DEL strain resulted in a significantly higher number of parasites and significantly fewer metacyclic forms than the NonDEL strain (**Figure 5**D). The results of the six assayed samples exposes the complex interplay between parasite- vector genetic variances, which are present in the local, as part of specific transmission cycles.

### The parasites carrying the sub-chromosomal deletion showed a reduced ability to escape killing from neutrophil NETs and infect macrophages, exposing variable infection outcomes in the mouse model

The ecto-3’-nucleotidase activity, which is absent in DEL strains, is involved in the infection process of *Leishmania* in macrophages^10,12^ and contributes to the escape of the parasite from neutrophil extracellular traps (NETs)^12,13^. Based on these observations, we performed a killing assay to compare the ability of DEL and NonDEL parasites to survive exposure to NETs. Results from the three pairs of strains selected from different geographic regions reveal that DEL parasites are less resistant to NETs (44.6, 62.5 and 75.6% survival) than NonDEL (78.8, 97.7, and 93.7%), relative to their controls (100%) (**Figure 6**A-C). In the macrophage infection assay, the percentage of infected cells by DEL parasites was lower than that by the Non-DEL at 24 hours (26.8 vs 48.6%; 32.2 vs 43.3%; 31.4 vs 44.2%) and at 48 hours (20.4 vs 35.8%; 21.6 vs 35.4%; 27.1 vs 38.5%;) (**Figure 6**D-F). No difference in the number of parasites per macrophage was observed, (**Figure 6**G-I), suggesting differences in parasite virulence are more relevant in the initial stages of infection, and less evident after *Leishmania* achieves the intracellular stage.

**Figure 6.**
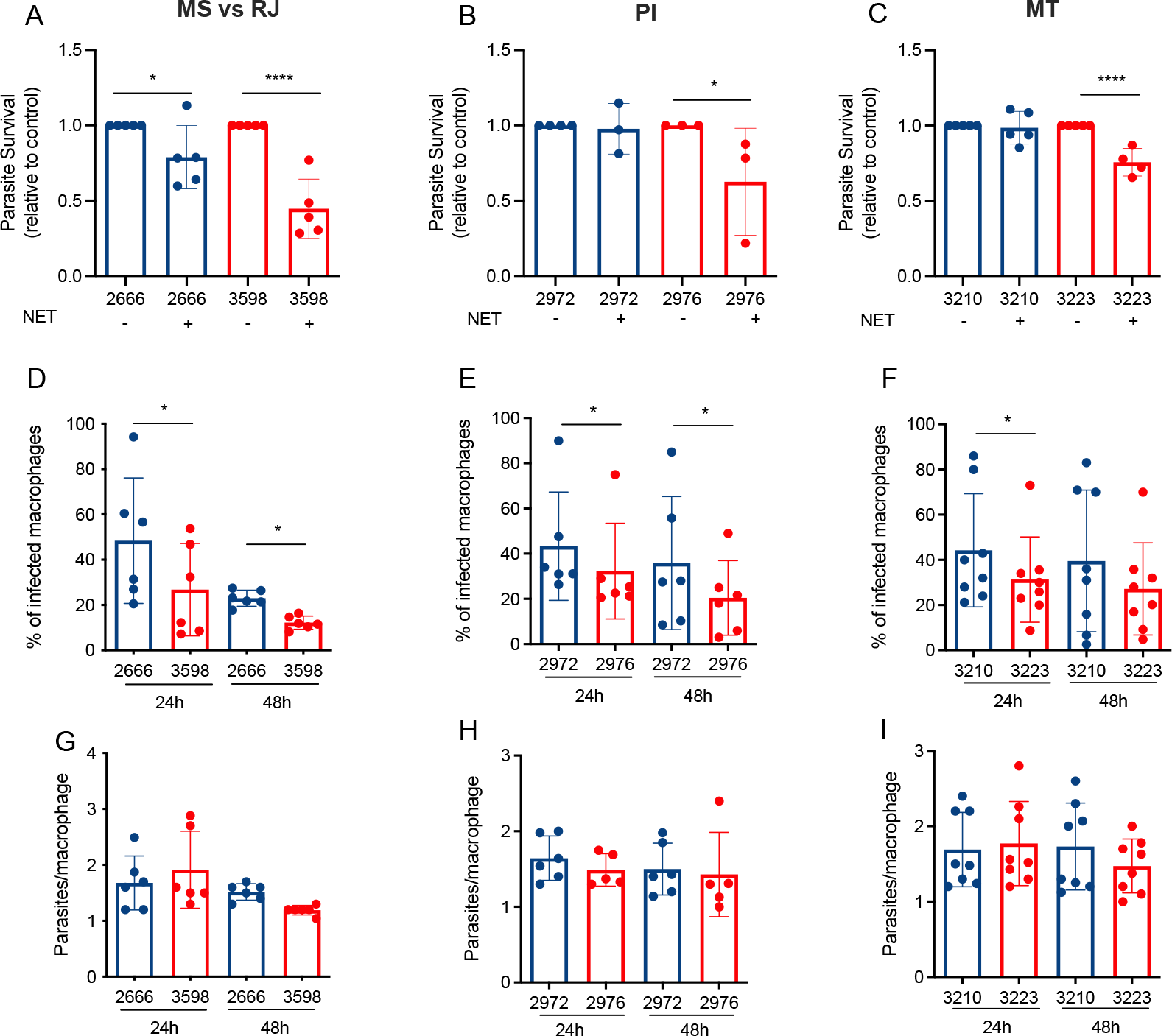
DEL parasites have a reduced ability to infect macrophages and escape killing from neutrophil NETs. (A-C) L. infantum metacyclics [5x10^5^] were incubated in the presence or absence of supernatants enriched in NETs for 4h at 35°C. Alamar blue (10% v/v) was added, and cells were incubated for more 4h at 35°C. Alamar blue fluorescence was read at 540/590 nm excitation/emission on a SpectraMax fluorimeter. Results are expressed as relative to control (parasites without NETs). Results of at least 3 independent experiments are shown as mean ± SEM. One-way ANOVA followed by Fisher’s LSD post-hoc comparison tests was performed. *p<0.05 and ****p<0.01. (D-F) Raw macrophages (2x10^5^) were seeded onto coverslips and then infected with Leishmania parasites at a cell ratio of 1 macrophage to 5 parasites. After 24h at 35°C, free parasites were washed out and coverslips were either fixed or incubated with medium (RPMI + 2.5% FBS) for more 24h (48h time point). Cells were stained with Panoptic dye kit and the number of infected macrophages (D-F) and the number of amastigotes/macrophage (G-I) were counted. Results are shown as mean ± SEM. Wilcoxon t-test was performed. *p<0.05. Blue bars = NonDEL; red bars = DEL strains.

To further investigate the phenotypic variances and the effects during the early stages of infection, the ear model assay in BALB/c mice was applied to assess the virulence of DEL/NonDEL strains from the previous experiments. Metacyclic parasites (10^5^) from passage-controlled cultures were inoculated intradermally at the inner ear and, 12-15 hours post-infection, neutrophil and monocyte recruitment were determined by flow cytometry. The data show reduced neutrophil and monocyte recruitment for DEL strains from the RJ vs MS pair (1.4 vs 2.1% of CD11b+ cells, respectively) (**Figure 7**A-B). The draining lymph nodes were harvested for parasite quantitation at the same time point (**Figure 7**C) exhibiting higher parasite load for the DEL strains from RJ (2.2 vs 0.5) and MT (4.6 vs 1.5 eq. par/mg tissue).

**Figure 7.**
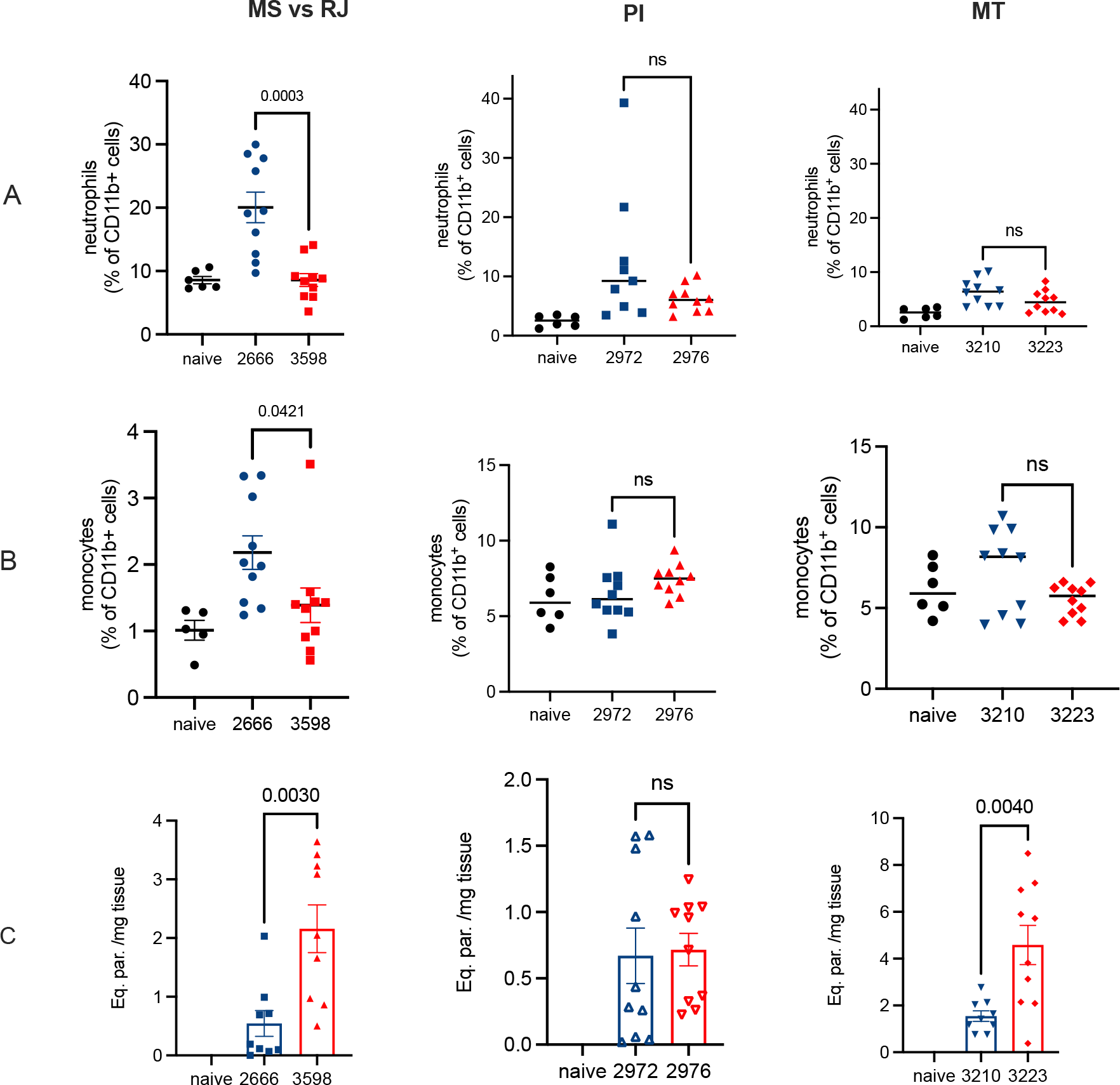
Neutrophil and monocyte recruitment to the site of infection by DEL and NonDEL parasites and parasite load from the draining lymph nodes. A - B) percentage of CD11b+ cells obtained from inoculated ear 12-15hours PI (neutrophil and monocyte recruitment, respectively). Blue= NonDEL strains; Red= DEL strains. L. infantum metacyclic (10^5^) was inoculated in the inner ear of anesthetized BALB/c mice. 12-15 hours post-infection animals were euthanized, and ears were collected in RPMI. Ordinary one-way ANOVA. C) Parasite load expressed by equivalent parasite per mg of the draining lymph nodes from each ear. Material was collected and preserved in DNAShield for further DNA isolation and qPCR. Strains used in the assays were geographically unrelated: RJ= Rio de Janeiro; MS= Mato Grosso do Sul; MT=Mato Grosso; PI=Piauí.

## Discussion

The presence of a 12 Kb deletion on chromosome 31, exclusively found in American *L. infantum* strains, is both highly prevalent and widely distributed geographically^4^. These epidemiological characteristics imply that the deletion provides a fitness advantage to this population of parasites under New World conditions, despite its adverse effect on the parasite’s crucial purine salvage pathways due to the depletion of 3’NU/NT activity. Compensatory strategies and potential associated epidemiological consequences in the American Visceral Leishmaniasis (AVL) scenario due to the circulation of the distinct genotypes is a critical open question. On this ground, a first and pivotal action is to investigate the more prominent biological consequences of the deletion and assess its effect on parasite / hosts interaction.

In this study, we utilized biological replicates of geographically unrelated *L. infantum* mutant strains to explore the substantial differences between DEL and NonDEL phenotypes and the intricate balance between fitness gain and cost inside the mammalian and insect hosts. Our in vitro data corroborates previous findings on the reduced or absent 3ʹ NT activity of geographically unrelated DEL promastigotes^4^. Furthermore, these DEL parasites exhibited, on average, higher rates of in vitro metacyclogenesis, likely triggered by nutritional stress resulting from the compromised purine salvage route dependent on 3’NU/NT activity^5^. The deficiency in 3’NT activity appears to induce a state akin to “starvation,” prompting an increase in the expression of NT1, a membrane transporter associated with purine uptake. This corroborates observations in *L. donovani* parasites deprived of purines, in which rapid amplification of the NT1 adenosine/pyrimidine nucleoside transporter gene was observed^8^, suggesting this as an alternative adaptive compensatory mechanism. Further experimentation under controlled purine-deprived conditions would offer deeper insights into the biological behavior of DEL parasites. It’s also worth considering the composition of vector saliva as a potential molecular modulator of DEL and NonDEL strain behavior during blood feeding. Previous studies have reported the presence of AMP and purines (adenosine) in the saliva of *Phlebotomus papatasi* and *P. duboscqi* sand fly, but, curiously, not in *L. longipalpis*^28^. These differences could influence the behavior and interactions of these parasites in each sand fly species. Delving into this aspect further could provide valuable insights into the transmission dynamics of DEL and NonDEL strains.

We further noted that the 3’NT activity in NonDEL is high in promastigotes, particularly in metacyclic forms. This implies that the enzyme may play an important role during the intravectorial phase or the early stages of mammalian infection. This raises the intriguing question of how DEL parasites, lacking 3’NU/NT, manage to overcome this enzyme deficiency and effectively infect hosts. It is conceivable that the higher rate of metacyclogenesis aids in overcoming this challenge and promotes successful infection. Additionally, our finding of higher transcripts of META2 in DEL metacyclic-enriched cultures suggests enhanced resistance to oxidative and heat shock stress^26^ for the infective form of these strains.

Increased metacyclogenesis leads to a lower parasite burden in the sand fly midgut, as this form is non-replicative. However, at the population level, it enhances transmissibility by the sand flies, thereby promoting geographic spread. In our experiments involving the main vector of *L. infantum* in the Americas, *Lutzomyia longipalpis*, a stronger signal was evident: two out of three DEL strains showed both a higher ability to colonize the insect’s midgut and a tendency towards increased metacyclogenesis. Increased parasite load in the vector midgut implies a higher dose of infective inoculum and may thus enable the parasite to establish infections ^29,30^. These findings suggest that the deletion enhances fitness during the vector-borne life cycle stage in epidemiologically significant sand fly species. Thus, considering the pivotal role of *L. longipalpis* in the transmission cycle of AVL, it is probable that the interaction between DEL parasites and this vector species facilitates the spread of this genotype in the Americas. However, the extent of this phenomenon may vary depending on the geographical region and the specific population of *L. longipalpis* involved^31^.

Contrary to the results of the sand fly experiments, the DEL parasites exhibited reduced fitness in the *in vitro* assays. This was evidenced by their lower virulence in macrophages and the diminished ability to escape killing by NETs. The results corroborate reports demonstrating that the downregulation of ecto-nuleotidases leads to decreased infectiousness^9^ and lower number of surviving parasites following exposure to NETs^12^. In parallel, DEL parasites induced reduced or similar-to NonDEL effect on neutrophil and monocyte recruitment in the mouse ear infection assay. These experimental findings of decreased infectiousness conflict with epidemiological data reporting a high prevalence of DEL parasites associated with human and canine cases of VL in the Americas^4,15^. Thus, within the complex and not yet fully understood course of VL pathogenesis, it is plausible that mutant parasites cause effective yet less pathogenic infection compared to NonDEL strains. Such variations in infectiousness and pathogenicity relate to host-parasite strategies of disease tolerance or resistance^32^. Each of these strategies might have distinct evolutionary outcomes: resistance could reduce parasite prevalence, whereas tolerance could be neutral towards, or increase, prevalence in the population, ultimately reaching epidemiological consequences^32^. It is conceivable, thus, that the emergence of this potentially virulence-attenuating trait among New World *L. infantum* prompts disease tolerance/accommodation mechanisms within hosts, thereby favoring transmission throughout the DEL parasite’s life cycle^33^. Over time, such traits would likely undergo positive selection, especially if these mutant strains evolve strategies for long-term persistence within the host^34^. This scenario is better recognized in *Toxoplasma gondii*^35^ in which increased tolerance by the host benefits the parasites, but it has not yet been fully elucidated in *Leishmania*.

The DEL mutant strains, characterized by reduced pathogenicity, may exploit the trade-off between cost and benefit in immunity. They can infect and replicate, possibly without inducing exacerbated clinical signs that prompt treatment or culling of the main urban reservoir, the domestic dog, which is an undesirable consequence for parasite transmission. If this holds true, it strengthens the importance of detecting asymptomatic individuals as a policy to control the spread of disease. Animals without clinical signs would remain undetected for longer periods of time, representing a continuous source of infection for the sand flies in the area, favoring parasite spread^36,37^. Simultaneously, these DEL strains possess a remarkable ability to colonize the vector and undergo metacyclogenesis, generating a highly effective inoculum. Taken together, these attributes signify a gain in fitness at the population level and help elucidate the higher prevalence of deleted parasites in Brazil. This scenario, coupled with the observed natural resistance of DEL parasites to MIL, highlights the epidemiological and clinical significance of identifying the genotype of the infecting *L. infantum* in both animals and humans.

## Methods

### Parasites origin, maintenance and confirmation of species and genotype

All *L. infantum* strains sequenced in this study were obtained from the Coleção de Leishmania da Fundação Oswaldo Cruz (CLIOC). Associate information to the strains selected is depicted in Figure 1 from supp material. In all cases, *Leishmania* was isolated from human or dog patients as part of normal diagnosis and treatment with no unnecessary invasive procedures and with written and/or verbal consent recorded at the time of clinical examination. All strains were cultured in biphasic medium (Novy–MacNeal–Nicolle (NNN) + Schneider’s added Fetal Calf Serum 20%) prior to genomic DNA extraction (DNeasy Blood & Tissue Kit, Qiagen). Species were confirmed by multilocus enzyme electrophoresis (MLEE) as presented in the internal SOP. Real time qPCR protocol previously published^4^ was applied to detect and/or confirm the deleted genotypes. Two different batches of cryopreservation were prepared: culture-adapted parasites (> than 20 passages), and passage-controlled parasites isolated from BALB/c mouse spleen and/or liver to be used in specific assays.

### Metacyclic enrichment by PNA

Metacyclic enrichment was performed following protocols previously published^38^. Briefly, parasites were washed twice in 50 ml Schneider (pure) by centrifuging 15 min at 3000 X g, 4C, in a 50-ml, resuspended in Schneider (pure), counted, and brought to 1–2x 10^8^/ml. The concentration of 0.01 vol of 5 to 10 mg/ml PNA was added to a final concentration of 50 to 100 μg/ml and parasites were incubate at room temperature for 15 to 30 min. After centrifugation for 5 min at 200 x g, at 4°C the collected supernatant was washed twice and resuspend in small volume of PBS for counting. The desired concentration can be obtained for inoculation preparation in PBS1x or for material to be preserved in TRizol for RNA isolation.

### pH/temperature stressed culture (“axenic amastigote”)

promastigotes were transferred from the conventional culture condition (Shneider plus FCS 20%) to pure Schneiderat pH6 and kept for 4 days at 37◦C. Morphology changes were monitored by light microscopy. In the fourth day, cells were subjected to 3’NT activity measurement.

### 3’NT activity measurement

The 3’-nucleotidase activity was determined as previously described^11,12^ for NonDEL, HTZ and DEL strains. Briefly, the ecto-3’-nucleotidase activity was determined by measuring the rate of inorganic phosphate (Pi) production as previously described^10^. *L. infantum* parasites (2×10^7^ cells/mL) were incubated at 37 °C for 60 min in 0.5 mL of a reaction mixture containing 116mM NaCl, 5.4mM KCl, 5.5mM D-glucose, 50mM HEPES buffer (pH 7.4) and 1mM 3’-AMP as the substrate. Reactions were started by adding the cell suspension and stopped by centrifugation (1500×g for 15 min), and 0.1 mL of the supernatant was added to 0.1 mL of Fiske-Subbarow reactive mixture to measure the Pi released into the supernatants at 650 nm. The concentration of Pi was determined using a standard curve for comparison. The ecto-3’-nucleotidase activity was calculated by subtracting the nonspecific 3’-AMP hydrolysis measured in the absence of cells. Experiments were performed at least twice, with similar results obtained from at least three separate cell suspensions. Values obtained for each strain was normalized by the ecto-3’-nucleotidase activity of *L. amazonensis* and unpaired t-test applied using PRIM 9. For the comparison between DEL, NonDEL and HTZ groups, ANOVA was applied and *p<0.05 was considered as statistically different.

### DNA isolation, standard curve and quantitative Real time PCR (qPCR)

Absolute quantitation of parasites in DNA isolated from lymph nodes was obtained using Standard curves built with 1:10 serially-diluted *Leishmania* DNA, ranging from 10^7^ to 10^1^ parasites per 20 milligrams of tissue from non-infected mice. The protocol includes the TaqMan™ system (Applied Biosystems® CA, USA) with mouse GAPDH (glyceraldehyde-3-phosphate dehydrogenase) as endogenous control (host target) (VIC™/MGB probe) and the 18S rDNA (FAM™/MGB probe) target for *Leishmania* (Fw 5’-GTACTGGGGCGTCAGAGGT-3’; 18S rDNA Rv 5’- TGGGTGTCATCGTTTGCAG-3’ and the probe 18S rDNA Tq 5’- FAM-AATTCTTAGACCGCACCAAG-NFQ-MGB-3’). The assay was performed, according to the following conditions: 95 ◦C for 10 min, 95 ◦C for 15 s and 59 ◦C for 1 min, 45 cycles, qPCRs were carried out with Applied Biosystems VIA7 equipment from the *Plataforma de Análises Moleculares* (RPT09J) (Rede de Plataformas Tecnológicas Fiocruz). A DNA-free master mix as a No Template Control (NTC) and all samples and controls were performed in duplicate.

### RNA extraction, cDNA synthesis and relative quantitation by Real Time qPCR

Total RNA was isolated using TRIzol (Invitrogen) following the available protocol (Sigma-Aldrich). cDNA was synthetized with the SuperScript IV Synthesis Kit (Invitrogen) as recommended by the manufacturer. Both, total RNA and cDNA were normalized after quantification in NanoDrop 2.000 (Thermo Scientific). Relative quantitation of transcripts was done in a Quant Studio Real-Time equipment, from the *Plataforma de Análises Moleculares* (RPT09J) (*Rede de Plataformas Tecnológicas Fiocruz*). Reaction conditions and primers used were as follows: 0.3 micromolar of each primer and 1x Syber Green (Applied Biosystems). 15 sec at 95 ◦C; 1 min annealing and extension at 60◦C, for 40 cycles. Alfa-tubulin was used as reference gene for the Delta-Delta Ct method. Primers are depicted within Supp. Material. The NonDEL strain IOCL 3124, from Portugal, was used as a calibrator. Fold change comparison between DEL and NonDEL groups was performed by unpaired Mann Whitney Rank Sum in PRISM 9. *p<0.05 was considered as statistically different.

### Neutrophil purification

Human neutrophils from the peripheral blood of healthy blood donors were isolated by density gradient centrifugation (Histopaque; Sigma–Aldrich) followed by hypotonic lysis of erythrocytes, as described previously^39^ .Purified neutrophils (≥95% of the cells) were resuspended in RPMI medium 1640 (Sigma). All experiments involving neutrophils were performed in RPMI. All procedures involving human blood were performed in accordance with the guidelines of the Research Ethics Committee (Hospital Universitário Clementino Fraga Filho, UFRJ, Brazil), protocol number: 4261015400005257

### Visualization of NETs

Neutrophils (1 × 10^5^) were allowed to seed on 0.001% poly-L-lysine-coated coverslips and then incubated with promastigotes of *L. infantum* 2666 or 3598 (1 × 10^5^). After 3h, slides were fixed with 4% paraformaldehyde and stained with DAPI (10 ug/mL; Sigma) for 10 min. Images were taken on an EVOS® FL Cell Imaging System.

### Production of NETs-enriched supernatant and parasite survival assay

Neutrophils (8 × 10^6^) were activated with PMA (100 nM) for 4 h at 37 °C/5% CO2. Supernatants were recovered and the quantification of NETs was performed with the Picogreen dsDNA kit (Invitrogen, Life Technologies) as described^39^. *L. infantum* metacyclics [5x10^5^] were incubated in the presence or absence of supernatants enriched in NETs (500 ng/mL DNA) supplemented with 1%FBS for 4h at 35°C/5% CO2. Alamar blue fluorescence was read at 540/590 nm excitation/emission on a SpectraMax fluorimeter (Molecular Devices), using an excitation of 540 nm and emission of 600 nm. Data were normalized based on the fluorescence of control parasites cultured in the absence of NETs. Wilcoxon t-test analysis was performed and *p<0.05 was considered as statistically different.

### RAW 264.7 cell line infection

The RAW 264.7 murine macrophage cell line was maintained in RPMI 1640 media containing 10% heat-inactivated FBS. Raw macrophages (2x10^5^) were seeded onto coverslips and then infected with *Leishmania* parasites at a cell ratio of 1 macrophage to 5 parasites. After 24h at 35°C, free parasites were washed out and coverslips were either fixed or incubated with medium (RPMI + 2.5% FBS) for more 24h. Cells were stained with a Panoptic dye kit and the number of infected macrophages, and the number of amastigotes/macrophages were counted. Wilcoxon t-test analysis was performed and *p<0.05 was considered statistically different.

### Experimental infections of sand flies

Experiments were performed using sand flies from a laboratory colony of *L. longipalpis* established from sand flies caught in Jacobina (Bahia, Brazil), *L. migonei* caught in Baturité (Ceará, Brazil), and *P. perniciosus* (Spain) using standard methods^40^. The insects were fed on 50% sucrose *ad libitum* and blood-fed on heparinized rabbit blood when needed. The insects were maintained at 27 ± 1 °C, humidity of 80–95%, and a photoperiod schedule of 12 h light/12 h dark. Three to five-day-old female sand flies were fed through chick skin membrane on heparinized rabbit blood containing 5×10^5^ *Leishmania* promastigotes/mL of each of the following strains at late log phase: IOCL 2666 (NonDEL), 3134 B1 (clone HTZ), IOCL 3598 (DEL), IOCL 3210 (non-DEL) and IOCL 3223 (DEL) isolated from Mato Grosso state (MT); and IOCL 2972 (non-DEL) and IOCL 2976 (DEL) isolated from Piauí state (PI). The midguts of insects were dissected 72 (day 3) and 192 hours (day 8) after infection. The number of infected sand flies was expressed as percentage of infected sand flies calculated in comparison of number of dissected guts. The number of parasites present per midgut was estimated from dissected guts homogenized in 50 μL of 0.85% NaCl + 1% formaldehyde solution, using a hemocytometer and plotted in log scale excluding zero values from non-infected sand flies. Localization of parasites in the sand fly gut was observed under 40x magnification^41^ and expressed as percentage of sand flies that developed stomodeal valve stage infection in comparison to the number of dissected insects. Morphology of parasites was determined from 5 to 8 sand fly gut smears on glass slides stained with Giemsa, photographed under 100x magnification light microscopy, and measured using ImageJ software^42^ and expressed as percentage of metacyclic promastigote forms in comparison to total number of analyzed parasite images from infected gut smears. A minimum of 3 independent experiments were preformed, and insects were analyzed individually. Statistical analysis was done using GraphPad Prism 6.07 computer software. Mann-Whitney test was used for pair wise comparisons of parasite numbers and the Chi-square test was used for contingency data of infected insects, infection localization, and parasite forms.

### Ear model infection in BALB/c mice

Animals were supplied by the Institute of Science and Technology in Biomodels (ICTB), Fiocruz and the experimental design was approved by Fiocruz Animal Welfare Committee (CEUA L-015/202-A2). Female BALB/c mice were anesthetized with xylazine and ketamine. Using a 30G ultra-fine needle, 10 µL of 10^5^ metacyclic-enriched culture in PBS were inoculated intradermally in the inner part of both ears. 13-16 hours after inoculation animals were euthanized with xylazine and ketamine and ears were collected in RPMI on ice for flow cytometry. Draining lymph nodes were harvested and preserved in a lysis solution from High Pure DNA Isolation Kit (Roche).

### Flow cytometry

Mice were euthanized, and the two sheets of ear dermis were isolated, placed in RPMI containing 0.2 μg/mL Liberase CI purified enzyme blend (Roche Diagnostics Corp.), and incubated for 1 h at 37 °C. After 1h, RPMI supplemented with FBS (10%) was added. Digested tissue was placed in a cell strainer (40 um; BD) and tissue debris were ground with the help of a syringe plunger. The resulting cells were stained with the Fixable Yellow Dead Cell Stain Kit (Invitrogen). Unspecific staining was blocked by incubating the ear cell suspension with mouse serum (10%). Cells were stained for Ly6C (BD Pharmingen™, Cat #553104, Clone AL-21, Lot #7067529; 1:200; FITC), Ly6G (BD Pharmingen™, Cat #551461, Clone 1A8, Lot #7068711; 1:800; PE), CD11b (BD Pharmingen™, Cat #552850, Clone M1/70, Lot #7033964; 1:800; PE-Cy7). Data were analyzed on a Fortessa flow cytometer (BD). Cells were acquired based on forward and side scatter, and live cells and data were analyzed with FlowJo Software 4.3.

## Funding

This work was supported by: PTR (Programmes Tranversaux de Recherche) from Institut Pasteur Paris [grant number PTR 425-21]; FIOCRUZ -PAEF [grant number IOC-023-FIO-18-2- 63]; Coordenação de Aperfeiçoamento de Pessoal de Nível Superior (CAPES) [grant number Finance Code 001]; Conselho Nacional de Desenvolvimento Científico e Tecnológico (CNPq) [grant number Research Fellow, 302622/2017-9, Process No. 312573/2020-0]; Fundação Carlos Chagas Filho de Amparo à Pesquisa do Estado do Rio de Janeiro (FAPERJ) [grant number CNE E26-202.569/2019, ColBio2020 E26-210.285/2021, FAPERJ Emergentes E-26/010.002168/2019, and CNE 26/200.487/2023]; Czech Ministry of Education and ERD funds [grant number CePaViP CZ.02.1.01/0.0/0.0/16_019/0000759].

## Acknowledgements

We thank Rede de Plataformas Tecnológicas da Fiocruz -Plataforma de Análises Moleculares (RPT09J) and Coleção de Leishmania da Fiocruz (CLIOC) for providing the *Leishmania infantum* strains. Artificial Intelligence was exclusively applied during the final stage of manuscript preparation for English language enhancement.

## Conflict of interest

Authors declare no conflict of interest.

## Notes

### Competing Interest Statement

The authors have declared no competing interest.

