## Supplemental Fig.1 and Table 1 for "Gene deletion as a possible strategy adopted by New World *Leishmania infantum* to maximize geographic dispersion"

**Parasites origin, maintenance and confirmation of species and genotype**

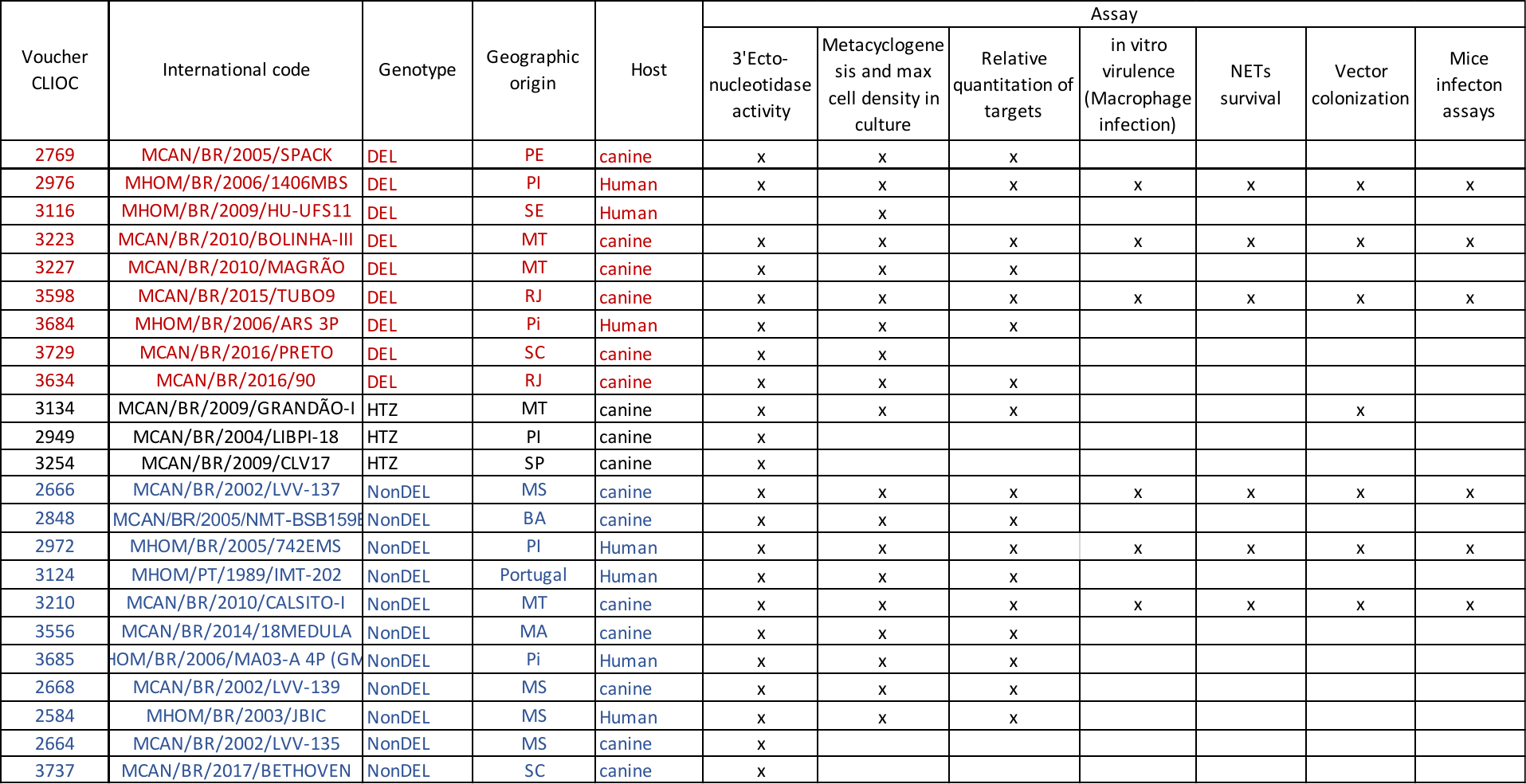

Figure 1. *L. infantum* strains selected for the assays, the respective genotypes, geographic origin, host, and assays applied. Genotype was determined by Whole Genome Sequencing (WGS) and further confirmed by qPCR. The geographic origin reflects the wide distribution of the selected samples (Northeast PE= Pernambuco, PI=Piaui, SE= Sergipe, BA= Bahia; Central-west: MT=Mato Grosso, MS= Mato Grosso do Sul; Southeast e South: RJ= Rio de Janeiro, SP= São Paulo, SC- Santa Catarina).

Initially, this set of 23 culture-adapted *L. infantum* strains, isolated from humans and dogs, comprising nine deletion-carrying (DEL), three heterozygous (HTZ), and eleven non-deleted (NonDEL) strains, were evaluated. Subsequently, subsets of strains were chosen for different assays, as outlined in Figure 1. Genotypes were identified via whole genome sequencing (WGS) and further confirmed by PCR. All *L. infantum* strains sequenced in this study were obtained from the Coleção de Leishmania da Fundação Oswaldo Cruz (CLIOC).

Table 1 . Primers used for gene expression assays.

| **Target** | **Primers** | **Primer design 3'-5'** |
| --- | --- | --- |
| *Nucleoside transporter 1* | NT1_F | CCGTGTCAAGTGGATGTTCG |
|  | NT1_R | TCGAGAACCACTTCGAGTCC |
| *Paraflagellar* | Paraflag_F | TACAACTGTGACCTGGCGAT |
|  | Paraflag_R | AAGGTCCTGGTTCGTCTTGT |
| *Amastin* | Amastin_F | ACGACCAGTGGAAGTTTTGC |
|  | Amastina_R | GCACAGCATAATGAAGCCGA |
| *Alfa-tubulin* | a-tubuln F | AGC ACA CCG ATG TTG CGA CGA T |
|  | a-tubulin R | GAT CAG GCG GTT CAC GTT CGT GT |
| *3'NT/NU* | 31_2380_Fw_Tq | GCTGAAGTCAGTGAGCATGGA |
|  | 31_2380_Rv_Tq | TTCTGATCGTAGTGGTTGTGCA |
| *META1* | META1 F | CGAGAGTGGCGAACATCATG |
|  | META1 R | ATCCCTGACTGAGAGCGTTC |
| *META2* | META2 F | AACGGTGGAAGCTTTGTCAC |
|  | META2 R | CTGCTTCGCTCTGTGTTGTT |
